# Expansion of the CRISPR toolbox in an animal with tRNA-flanked Cas9 and Cpf1 gRNAs

**DOI:** 10.1101/046417

**Authors:** Fillip Port, Simon L Bullock

## Abstract

We present vectors for producing multiple CRISPR gRNAs from a single RNA polymerase II or III transcript in *Drosophila*. The system, which is based on liberation of gRNAs by processing of flanking t †RNAs, permits highly efficient multiplexing of Cas9-based mutagenesis. We also demonstrate that the †RNA-gRNA system markedly increases the efficacy of conditional gene disruption by Cas9 and can promote editing by the recently discovered RNA-guided endonuclease Cpf1.

---

The clustered regularly-interspaced short palindromic repeats (CRISPR) system greatly facilitates genome engineering in many species (Doudna and Charpentier, 2014). The method involves site-specific DNA cleavage by a bacterial endonuclease - such as the prototypical CRISPR-associated 9 (Cas9) protein - which is directed to a specific genomic target site by a short, single guide RNA (gRNA). Imprecise repair of the DNA break can create loss-of-function insertion/deletion (indel) mutations, whereas homology-directed repair in the presence of an experimentally provided donor DNA can generate specific sequence alterations.

Despite the power of current CRISPR approaches, improvements are necessary to realise the full potential of this technology in basic research, as well as in applied settings. One current focus is the development of robust methods for simultaneous expression of several gRNAs in an organismal setting. Multiplexed gRNA expression would, for example, facilitate the generation of non-functional mutations in a single target gene, as well as the creation of models of polygenic human diseases. The use of multiple gRNAs is also expected to significantly reduce the risk of natural resistance to gene drives designed to combat insect-borne diseases such as malaria or Zika virus (Champer et al., 2016).

Multiplexing methods that liberate gRNAs from a single precursor RNA in the nucleus have the added advantage of allowing expression from RNA polymerase (pol) II promoters (Nissim et al., 2014), as opposed to the RNA pol III promoters that are conventionally used to avoid the modification of gRNAs and subsequent export to the cytoplasm. Coupling of gRNA production to RNA pol II transcription gives scope to regulate production of guide sequences in time and space, thus affording additional control of CRISPR mutagenesis. Strategies to excise multiple gRNAs from a single precursor transcript have been described (Nissim et al., 2014; Tsai et al., 2014; Xie et al., 2015). However, they have not yet been adopted for use in animal models.

Here we describe novel vectors for multiplexed gRNA expression in *Drosophila*. These plasmids are inspired by a tRNA-gRNA expression system recently described for genome engineering in rice (Xie et al., 2015), in which the endogenous tRNA processing machinery excises multiple functional gRNAs from a single precursor transcript in the nucleus (Fig. 1a). We first demonstrated that a flanking tRNA does not interfere with production of functional gRNAs in *Drosophila*. The *Drosophila* tRNA^Gly^ was inserted between the strong, ubiquitously active RNA pol III promoter *U6:3* and a previously characterised gRNA, which targets the eye pigmentation gene *sepia* (se) (Port et al., 2015). The *U6:3-tRNA-gRNA* (*U6:3-t::gRNA*) transgene was transformed into the *Drosophila* genome and combined through a genetic cross with an *act-cas9* transgene, which expresses Cas9 ubiquitously from the *actin5C* promoter (Port et al., 2014). The rate of germline transmission of loss-of-function *se* mutations by the *act-cas9/U6:3-t::gRNA* animals was similar to a control in which no tRNA was present in the gRNA expression construct (Fig. 1b). The presence of the tRNA in the construct also did not influence the rate of germline mutagenesis by a second gRNA, which targeted the pigmentation gene *ebony* (*e*) (Fig. 1b).

**Figure 1.**
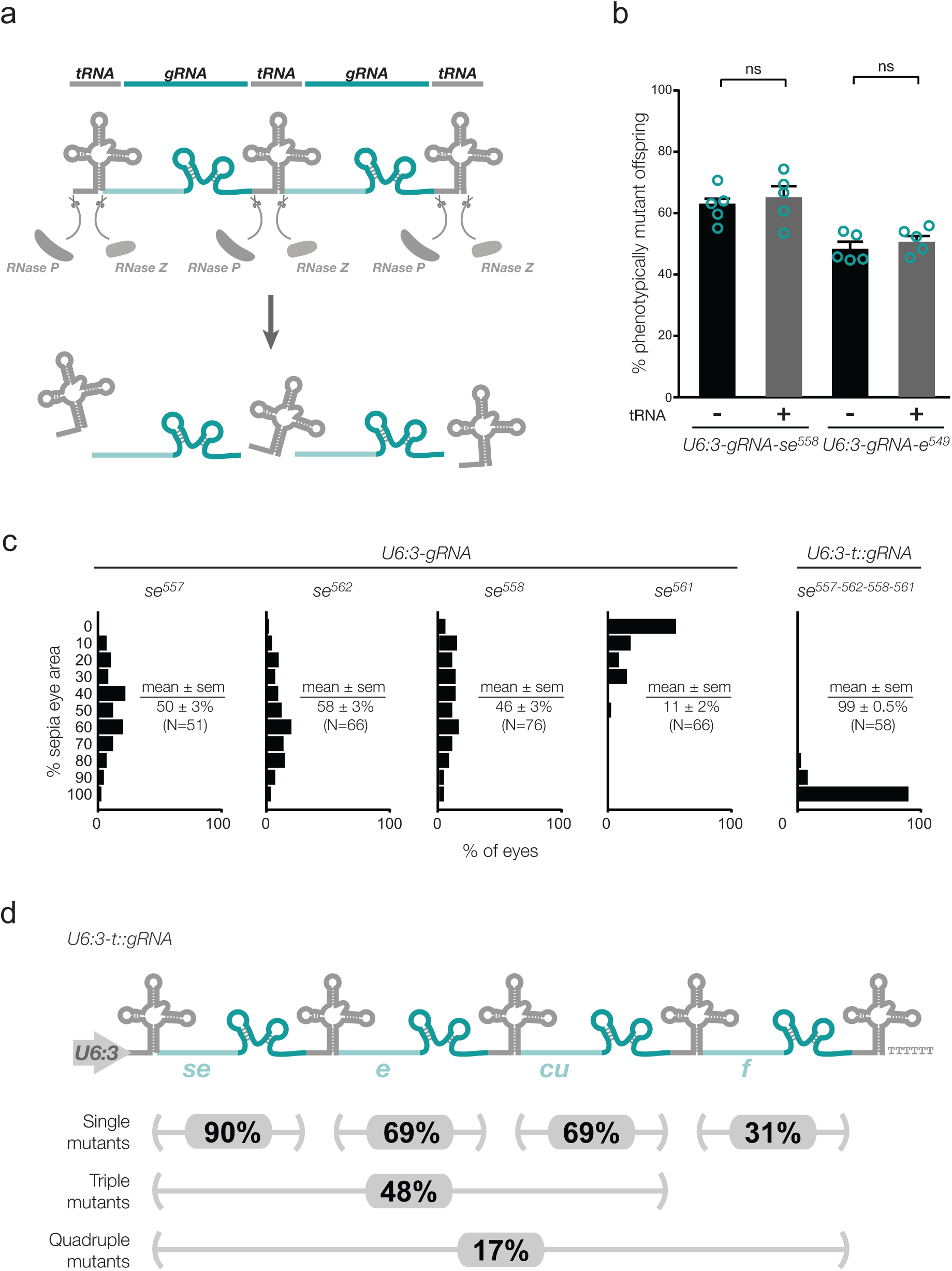
**Multiplexed Cas9 gRNA expression in *Drosophila* with the tRNA-gRNA expression system. (a)** Principle of the tRNA-gRNA system. gRNAs are excised from a long precursor transcript by the endogenous tRNA processing machinery (Xie et al., 2015). **(b)** Comparison of rates of target gene mutagenesis of gRNAs in the presence (-) or absence (+) of a single 5’ tRNA in the expression vector. *y*-axis represents rates of germline transmission of non-functional alleles of *se* or *e* by flies that are transgenic for both a*ct-cas9* and the indicated gRNA expression vector (five crosses per condition). ns, not significant; error bars, sem. **(c)** Comparison of rates of mutagenesis between individual gRNA transgenes targeting *se* and all four gRNAs expressed simultaneously from a *pCFD5 U6:3-t::gRNA* transgene. Data show proportions of eyes (10% bins) of *act-cas9/gRNA* animals that are sepia, which indicates non-functional mutations in both *se* alleles. N, number of eyes scored. **(d)** The use of *pCFD5* to generate mutations in multiple genes in a single step. Percentages of offspring of *act-cas9/U6:3-t:gRNA-se^588^-e^578^-cu^801^-f^801^* transgenic animals that inherit mutations in *se*, *e*, *cu* and *f* are shown. Mutation frequencies were determined by sequencing PCR products from the target loci in 48 offspring. The data therefore do not discriminate between functional and non-functional CRISPR alleles.

We next created *pCFD5*, a vector that facilitates cloning of several Cas9 gRNAs flanked by *Drosophila* or rice tRNA^Gly^ downstream of a single *U6:3* promoter (see Supplementary Methods for cloning protocol). Transgenic flies were generated that harbour a *pCFD5* construct containing four different gRNAs targeting *se*. We had previously shown that expression of the individual gRNAs in the presence of *act-cas9* results in biallelic disruption of *se* in only a fraction of eye tissue (Port et al., 2015). This was confirmed in the current study, with the mean amount of *se* mutant tissue per eye ranging from 11 to 58% for the single gRNAs (Fig. 1c). Incompletely penetrant mutagenesis with these gRNAs is due to either suboptimal cleavage by Cas9 or in-frame mutations at the target sites resulting in a functional protein product (Port et al., 2015). Strikingly, expressing all four gRNAs simultaneously with *pCFD5* in the presence of *act-cas9* resulted in biallelic disruption of *se* in 100% of eye tissue in almost all cases (Fig. 1c). Thus, using the tRNA-gRNA system to simultaneously express several gRNAs can circumvent the problems of functional in-frame mutations and gRNAs with low activity.

We also created transgenic flies harbouring a *pCFD5* construct containing four gRNAs that each target a different gene: *se*, *e*, *curled* (*cu*) and *forked* (*f*). Animals expressing both *act-cas9* and *U6:3-t::gRNA-se:e:cu:f* had a high penetrance of the visible phenotypes associated with biallelic disruption of each gene (Supplementary Fig. 1). Of the progeny of *act-cas9/U6:3-t::gRNA-se:e:cu:f* animals, 17% inherited indel mutations at all four loci (Fig. 1d). 48% of all progeny had mutations in *se*, *e*, and *cu*, which are located on the same chromosome. Creating a triple mutant chromosome by recombining three single mutations would be time consuming, especially for genes without a visible phenotype. Our results demonstrate that *pCFD5* allows efficient generation of complex genotypes in a short amount of time.

**Figure 2.**
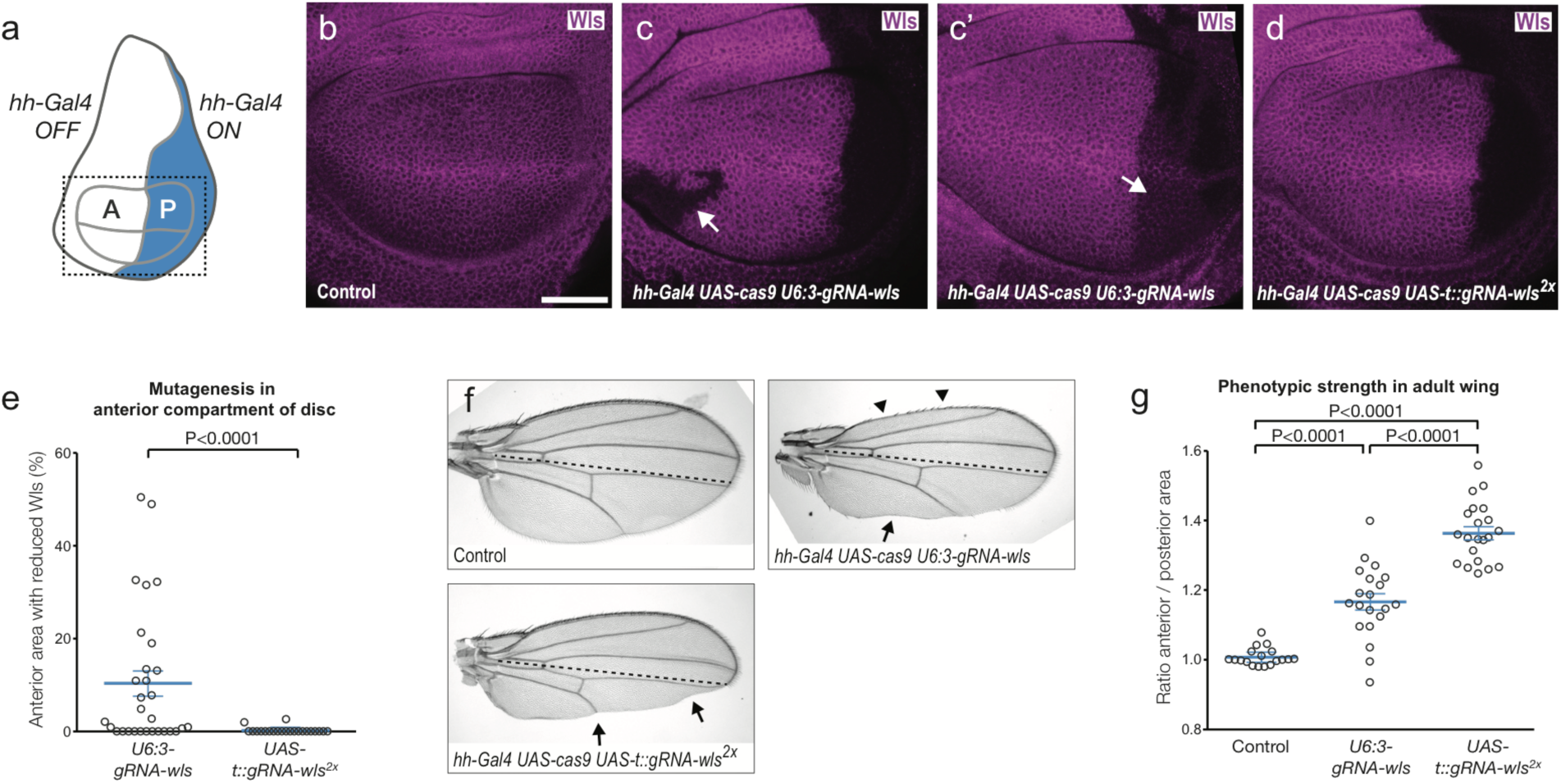
**The UAS-tRNA-gRNA system increases mutagenesis efficiency and tissue specificity of conditional CRISPR. (a)** Cartoon of a third-instar wing imaginal disc showing the expression pattern of *hh-Gal4*. Dotted rectangle shows area imaged. **(b** - **d)** Wing imaginal discs of indicated genotypes stained for endogenous Wls. **(b)** Wls is found throughout *hh-Gal4 UAS-cas9* control discs (control genotype is the same in **f** and **g**). Image is representative of 15 discs analysed. **(c, c’)** Introduction of a ubiquitously expressed *gRNA-wls* transgene results in incomplete mutagenesis in the posterior compartment and unwanted mutagenesis in the anterior compartment. **c** and **c’** show, respectively, examples of complete and partial loss of Wls immunoreactivity in the posterior compartment (arrow in **c’** indicates a large posterior clone that retains Ws signal). The disc in **c** also contains a large Wls-negative clone in the anterior compartment (arrow). Residual Ws staining in the posterior compartment was observed in 11/30 discs of this genotype. Reduced Wls staining in the anterior compartment was observed in 19/30 discs. **(d)** Example of completely penetrant disruption of Wls in the posterior compartment using a *UAS-t::gRNA* transgene. This phenotype was observed in 22/22 discs. The disc in **d** is also representative of the lack of targeting in the anterior compartment with this gRNA transgene (small clones with reduced Ws signal were observed in only 2/22 discs). **(e)** Quantification of ectopic mutagenesis in the anterior compartment in wing discs. **(f, g)** Comparison of adult wing phenotypes between *hh-Gal4 UAS-cas9 UAS-gRNA-wls^2x^* and *hh-Gal4 UAS-cas9 U6:3-gRNA-wls* animals. **f** shows representative examples of wing phenotypes. Arrows show loss of posterior wing margin tissue, resulting from absence of Ws in the posterior compartment of the wing imaginal disc. *UAS-gRNA-wls^2x^* leads to a greater loss of tissue at the posterior wing margin. Arrowheads show examples of loss of bristles at the anterior wing margin caused by targeting of Wls in the anterior compartment in the presence of *U6:3-gRNA-wls*. Dotted lines indicate the anterior-posterior boundary. **g** shows quantification of phenotypic strength in the anterior vs posterior compartment of the adult wing. In **e** and **g**, each circle represents one wing imaginal disc or one wing, respectively, and horizontal blue lines indicate mean ± sem. Statistical significance was evaluated with a Mann-Whitney test.

We next tested the performance of tRNA-gRNA vectors in another important application: the disruption of gene function in specific cell types in multicellular organisms. The use of CRISPR for such applications is desirable because, in principle, null mutations can be induced that produce more penetrant effects than RNAi knockdown approaches. Conditional CRISPR mutagenesis has been attempted in *Drosophila* by tissue specific expression of Cas9 with the binary, RNA pol II-based *UAS/Gal4* system in combination with individual gRNAs expressed from ubiquitously active RNA pol III promoters (Port et al., 2014; Xue et al., 2014). However, in our hands these approaches lead to poorly penetrant phenotypes in the target tissue or mutagenesis outside the desired domain (Port et al., 2014 and unpublished observations). Both these factors can complicate the interpretation of experimental findings significantly.

Based on our previous findings with *se* we reasoned that the expression of multiple guides from tRNA-gRNA vectors should increase the penetrance of loss-of-function phenotypes in the Gal4 expression domain. We also hypothesised that our approach would address the issue of mutagenesis outside the intended domain. Such mutagenesis is caused by stochastic expression of the *UAS-cas9* transgenes in the presence of the ubiquitously expressed gRNA. Expressing both Cas9 and gRNA under *UAS* control should increase tissue-specificity, because it is unlikely that sufficient ‘leaky’ expression of both transgenes would occur at the same time in the same cell. Previous attempts to express functional gRNAs from RNA pol II promoters in *Drosophila* using conventional vectors have been unsuccessful (Ren et al., 2013), presumably due to modification of the RNA and export to the cytoplasm. As described above, tRNA-mediated excision of gRNAs from larger transcripts in the nucleus should allow expression of active guides from this class of promoters.

We integrated into the genome a UAS-tRNA-gRNA construct (*UAS-t::gRNA-wls^2x^*), which contains two different Cas9 gRNAs targeting the ubiquitously expressed Wnt secretion factor *wntless* (*wls/evi*). We compared the activity of this transgene to *U6:3-gRNA-wls*, which has one of the *wls* gRNAs under the control of the *U6:3* promoter (Port et al., 2014). This was achieved by combining each transgene with an optimised *UAS-cas9* transgene (see Methods), and the *hedgehog-Gal4* (*hh-Gal4*) driver (Tanimoto et al., 2000), which activates expression of UAS transgenes in the posterior compartment of wing imaginal discs (Fig. 2a). Only 60% of *hh-Gal4 UAS-cas9 U6:3-gRNA-wls* wing discs had no detectable Wls protein in cells of the posterior compartment (n = 30; Fig. 2b – c’). In contrast, all *hh-Gal4 UAS-cas9 UAS-t::gRNA-wls^2x^* discs had lost Wls immunoreactivity throughout the posterior compartment (n = 22; Fig. 2d). Thus, the UAS-tRNA-gRNA construct induced much more efficient biallelic mutagenesis within the target domain.

Moreover, the *U6:3-gRNA-wls* transgene caused extensive mutagenesis outside the *hh-Gal4* expression domain. In 60% of the *hh-Gal4 UAS-cas9 U6:3-gRNA-wls* wing discs, there was anterior compartment tissue that had no (9 clones) or reduced levels (12 clones) of Wls protein (Fig. 2c, e). These patches ranged in size between a few cells to approximately half the size of the anterior compartment (Fig. 2e). Despite more efficient mutagenesis in the posterior compartment than observed with *U6:3-gRNA-wls*, only 10% of the *hh-Gal4 UAS-cas9 UAS-gRNA-wls^2x^* discs had patches of cells in the anterior compartment with reduced Wls levels, and these were very small (Fig. 2e). The effects we observed on disruption of Wls expression in the anterior and posterior compartment of the wing disc with *U6:3-gRNA-wls* and *UAS-t::gRNA-wls^2x^* were mirrored by the strength and location of *wls* mutant phenotypes in the adult wing (Fig. 2f, g). Together, these results demonstrate that *UAS-t::gRNA* expression vectors make it possible to perform conditional CRISPR mutagenesis with high efficiency and tight spatial control.

The data above demonstrated that the tRNA-gRNA system enhances Cas9-based CRISPR applications in *Drosophila*. Next, we explored whether this system can augment genome engineering by the RNA-guided endonuclease Cpf1. Cpf1 has a number of properties distinct from Cas9 that have the potential to broaden genome engineering applications (Zetsche et al., 2015). Because Cpf1 uses a poly-T protospacer adjacent motif (PAM) in the target DNA (Zetsche et al., 2015), as opposed to Cas9’s G-rich PAM (Jinek et al., 2012; Kleinstiver et al., 2015), the number of genomic sites for CRISPR is increased. Furthermore, the staggered cleavage of Cpf1 relatively far from the PAM (Fig. 3a) could conceivably facilitate CRISPR-induced homology-directed repair (Zetsche et al., 2015). However, genome engineering by Cpf1 had thus far only been demonstrated in cultured mammalian cells.

We constructed a *Drosophila* expression plasmid for production of AsCpf1 – one of the active Cpf1 variants in mammalian cells (Zetsche et al., 2015) – from the *actin5c* promoter. We also produced plasmids with a Cpf1 gRNA targeting *e* with or without flanking tRNAs under the control of the *U6:3* promoter. Co-injection of embryos with the *act-cpf1* plasmid and the *Cpf1 gRNA-e* plasmid without the tRNA resulted in transmission of non-functional *e* mutations to 1.6% of the progeny (Fig. 3b). Interestingly, the germline transmission rate was increased to 6% after injecting the *act-cpf1* plasmid and the plasmid encoding the *e* gRNA flanked by tRNAs (Fig. 3b). Targeted sequencing of the *e* locus confirmed the transmission of indel mutations at the gRNA target site (Fig. 3c). The vast majority of deletions were 10 – 20 bp in size (Fig. 3d). This is significantly larger than the deletions typically induced by Cas9, which rarely exceed 10 bp ((Moreno-Mateos et al., 2015); Fig. 3d). Analysis of more target sites will be necessary to confirm whether the creation of relatively large indels, which are more likely to be mutagenic, is a general feature of Cpf1-mediated mutagenesis.

**Figure 3.**
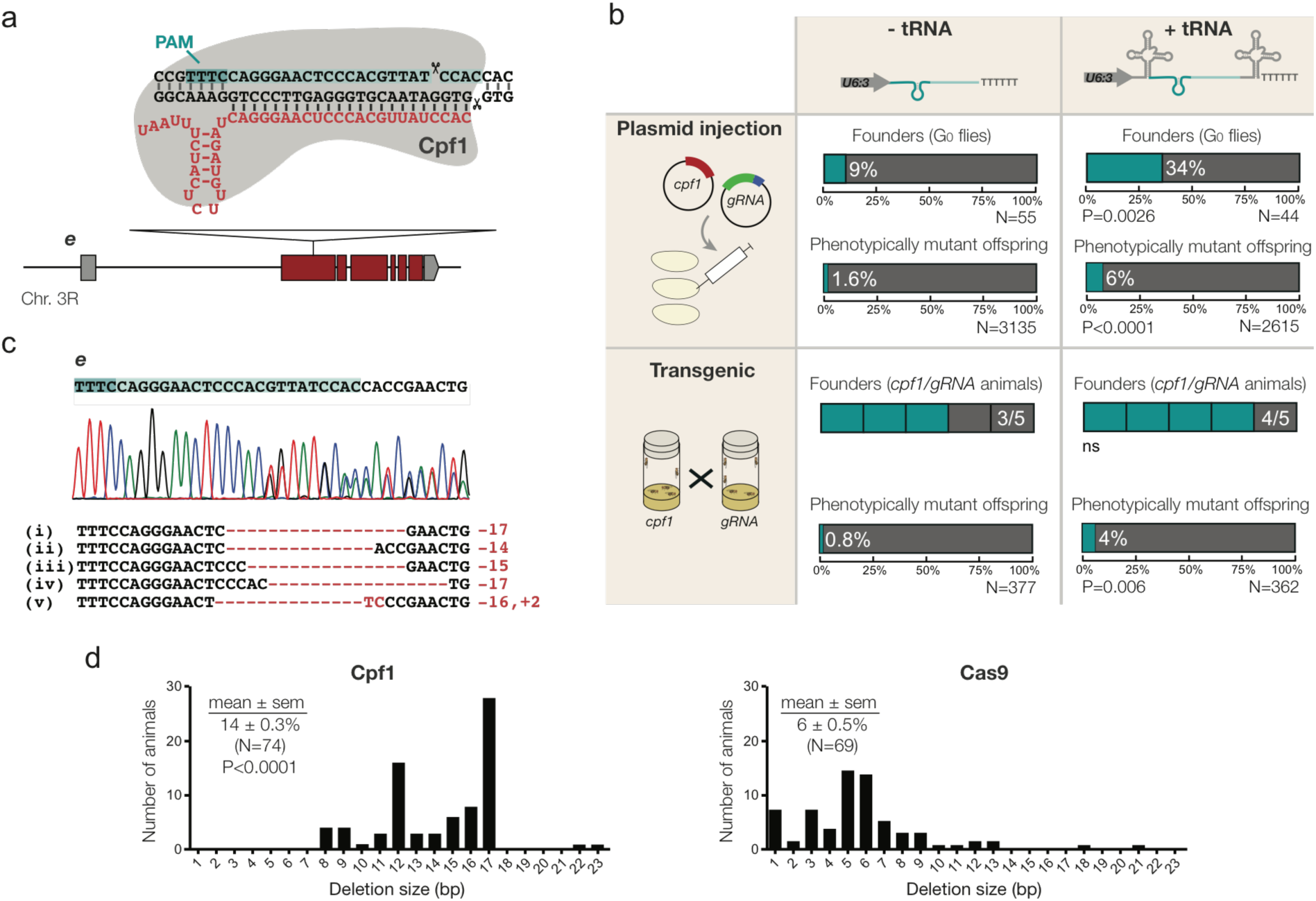
**Flanking tRNAs can enhance Cpf1 genome editing *in vivo*. (a)** Schematic of the Cpf1 target site in the *e* locus and the corresponding Cpf1 gRNA (red). Note that Cpf1 gRNAs are significantly shorter than Cas9 gRNAs and have an inverted topology (Zetsche et al., 2015). **(b)** Comparison of editing of *e* by Cpf1 in the presence or absence of flanking tRNAs in the gRNA expression vector. The experiment was performed either by co-injecting *act-Cpf1* (400 ng/µl) and *U6:3-gRNA* (200 ng/µl) plasmids into embryos or by providing both components as genomically integrated transgenes. A founder is a fly that transmits at least one non-functional *e* allele to the next generation. N, number of flies analysed. P values are for the comparison to the equivalent parameter in the absence of tRNA (two-tailed Fisher’s exact test). **(c)** Examples of Cpf1 -induced mutations in *e* that were transmitted through the germline. The chromatogram is from flies heterozygous for CRISPR allele iv. **(d)** Distribution of CRISPR-induced deletion sizes in *e* using Cpf1 and Cas9. Cas9 data are derived from *act-cas9 U6:3-gRNA-e* double transgenic flies (Port et al., 2014). Statistical significance (compared to the size of Cas9-induced deletions) was evaluated with a Mann-Whitney test. N, number of animals.

To test if increased mutagenesis with tRNA-flanked guides is a consistent feature of mutagenesis by Cpf1 in *Drosophila*, we generated flies with stable genomic integrations of *act-cpf1* and the *U6:3-gRNA* plasmids targeting *e*. We again observed a significant increase in germline transmission rates with the tRNA-flanked gRNA transgene compared to a transgene expressing the gRNA alone (Fig. 3b). Surprisingly, the germline transmission rates were similar for the same gRNA when the CRISPR components were provided by transgenes or by injected plasmids (Fig. 3b). This contrasts to the situation for Cas9 gRNAs, where mutagenesis is significantly increased by supply of the endonuclease and gRNA by transgenes (Kondo and Ueda, 2013; Port et al., 2014). This method results in the majority of alleles being mutagenised by Cas9 using the vast majority of gRNAs (Port et al., 2015). These observations suggested that Cpf1 might not perform as efficiently as Cas9 in *Drosophila* genome engineering experiments. Consistent with this notion, in plasmid injection experiments we did not detect any germline transmission of non-functional mutations using *act-cpf1* and three other Cpf1 gRNAs (targeting either *se* or the pigmentation gene *yellow* (*y*)), even in the presence of flanking tRNAs (Supplementary Fig. 2). We conclude that Cpf1 is able to act as an RNA-guided endonuclease in *Drosophila*. However, improvements will be necessary to transform Cpf1 into a robust genome editing tool. Our data indicating that flanking active Cpf1 gRNAs with tRNAs can enhance genome editing *in vivo* is a first step in this direction. Future experiments will investigate the mechanism by which tRNAs can increase the activity of Cpf1 gRNAs.

In summary, our data demonstrate that the tRNA-gRNA system can enhance Cas9 and Cpf1-mediated CRISPR mutagenesis in an animal model. In addition to the applications discussed above, we envisage that multiplexed gRNA expression with this method will prove valuable *in vivo* for disrupting the function of non-coding regulatory elements and for robust transcriptional activation (CRISPRa) and repression (CRISPRi), which often requires the action of multiple guides (Lin et al., 2015; Maeder et al., 2013; Mali et al., 2013a; Perez-Pinera et al., 2013). Furthermore, the use of flanking tRNAs will increase the repertoire of genomic sequences that can be targeted with novel high-fidelity Cas9 variants (Kleinstiver et al., 2016; Slaymaker et al., 2015). Because these endonucleases are unable to tolerate mismatches between the gRNA and genomic DNA, target site selection has been constrained by the need for a 5’ G in the guide sequence for optimal transcriptional initiation by RNA pol III. This is not the case when using tRNA-gRNA vectors with this polymerase, as transcription is initiated before the 5’ tRNA.

## METHODS

Additional information and updates are available from our website www.crisprflydesign.org.

### gRNA expression plasmids

Unless noted otherwise, PCRs were performed with the Q5 Hot-start 2x master mix (New England Biolabs (NEB)) and cloning was performed using the Gibson Assembly 2x Master Mix (NEB) following the manufacturer’s instructions. The sequence of each insert was verified by Sanger sequencing. gRNA sequences and other details of gRNA expression plasmids are presented in Supplementary Table 1.

#### pCFD5 (U6:3-(t::gRNA^cas9^)_1-6_)

The plasmid *pCFD3* (Port et al., 2014) was cut with BbsI and gel purified using standard procedures. A gBlock was ordered from Integrated DNA Technologies (IDT) that contained *Drosophila* tRNA^Gly^, a spacer containing two BbsI sites, a gRNA core sequence, a rice tRNA^Gly^ and appropriate homology arms (Supplementary Fig. 3). The gBlock was assembled with the *pCFD3* backbone by a Gibson Assembly reaction. Rice tRNA^Gly^ was chosen as the downstream tRNA since in preliminary experiments a construct with *Drosophila* tRNA^His^ at this position had reduced activity of 3’ gRNAs. A cloning protocol to generate *pCFD5* plasmids encoding one to six tRNA-flanked gRNAs is provided in the Supplementary Methods. *pCFD5* is available from Addgene (Plasmid 73914).

#### pUAS-t::gRNA-wls^2x^

Two tRNA-flanked *wls* gRNAs were cloned into the *pBID-UASC* backbone (gift from Brian McCabe (Wang et al., 2012), Addgene 35200). *pBID-UASC* was digested with EcoRI and XbaI and the linear plasmid gel purified. Inserts were generated by PCR using *pCFD5* as the template and the following primers: UAStRNAfwd1 *TGAATCACAAGACGCATACCAAACGA ATTCGGGCTTTGAGTGTGTGTAGACA* with UAStRNArev1 *CTCAGGTTCTCCAGTATGGTTGCATCGGCCGGGAATCGAACC* to generate a fragment encoding *Drosophila tRNA^Gly^*; UAStRNAfwd2 *ACCATACTGGAGAACCTGAGGTTTTAGAGCTAGAAATAGCAAG* and UAStRNArev2 *TGGCGAATATCACTTAGCAGTGCACCAGCCGGGAATCGAACCCGG* to generate a fragment encoding the first *gRNA-wls* and rice *tRNA^Gly^*; and UAStRNAfwd3 *CTGCTAAGTGATATTCGCCAGTTTTAGAGCTAGAA ATAGCAAG* and UAStRNArev3 *ACAGAAGTAAGGTTCCTTCACAAAGATCCTCTAGATGCACCAGCCGGGAATCGAACCCGG* to generate a fragment encoding the second *gRNA-wls* and rice *tRNA^Gly^*. The three inserts and the *pBID-UASC* backbone were assembled by Gibson Assembly.

#### pCFD6

To streamline cloning of gRNAs into a *pUAS- tRNA* vector, *pUAS-t::gRNA-wls^2x^* was modified to contain a BbsI cloning cassette. *pUAS-t::gRNA-wls^2x^* was cut with EcoRI and XbaI and the linear backbone gel purified. Inserts were generated by PCR using the following primers and templates: pCFD6fwd1: *GCCAACTTTGAATCACAAGACGC* and pCFD6rev1: *AGACCCTGCATCGGCCGGGAATCGAACCCG* with template *pUAS-t::gRNA-wls^2x^*; pCFD6fwd2: *CGGGTTCGATTCCCGGCCGATGC* and pCFD6rev2: *GCTATTTCTAGCTCTAAAACAGGTCTTCTGCACCAGCCGGGAATCGAA* with template *pCFD5*.*2* (where the first gRNA core sequence is modified according to (Dang et al., 2015)); pCFD6fwd3: *GAAGACCTGTTTTAGAGCTAGAAATAGCAAGT* and pCFD6rev3: *ACACCACAGAAGTAAGGTTCCTTC* with template *pUAS-t::gRNA-wls^2x^*. The three inserts and the *pBID-UASC* backbone were assembled by Gibson Assembly. In a second step, the three BbsI sites present in the vector backbone (in the *white^+^* marker) were removed by site-directed mutagenesis and by exchanging the *SV40 3’UTR* with the *p10 3’UTR* present in plasmid *pJFRC81* (gift from Gerald Rubin (Pfeiffer et al., 2012), Addgene 36432). Plasmids *pCFD6* and *pUAS-t::gRNA-wls^2x^* should become available from Addgene during April or May 2016.

#### pU6:3-hAsCpf1-gRNA

Expression plasmids for Cpf1 gRNAs were generated by Gibson Assembly using *pCFD5* digested with BbsI and XbaI and inserts generated by PCR using the following primers and *pCFD5* as a template: Cpf1gRNAPcr1fwd *TCGATTCCCGGCCGATGCATAATTTCTACTCTTGT AGAT-N23*(*target*)*-AACAAAGCACCAGTGGTCTAG* and Cpf1gRNAPcr1rev *TGCACCAGCCGGGAATCGAAC* to generate a fragment encoding the Cpf1gRNA and a rice tRNA^Gly^; and Cpf1gRNAPcr2fwd *ACAGACCCGGGTTCGATTCCCGGCTGGTGCATTTTTTGCCTACCTGGAGCCTGAGA* and Cpf1gRNAPcr2rev *TATACTGTTGCCGAGCACAATTGTCTAGAATGCATACGCATTAAGCGAAC* to generate a fragment encoding the genomic U6:3 terminator sequence directly downstream of rice tRNA^Gly^.

#### pCFD7

To make cloning of Cpf1 gRNAs more simple, we constructed *pCFD7*, which contains a BbsI cassette in place of the gRNA target site in *pU6:3- hAsCpf1-gRNA*. *pCFD7* was generated by Gibson Assembly using *pCFD5* digested with BbsI and XbaI and inserts generated by PCR using the following primers and *pCFD5* as a template: Cpf1gRNABbsIfwd *TCGATTCCCGGCCGATGCATAATTTCTACTCTTGTAGATGGGTCTTCGTTATCAGAAGACCTAACAAAGCACCAGTGGTCTAG* and Cpf1gRNAPcr1rev *TGCACCAGCCGGGAATCGAAC* to generate a fragment encoding the Cpf1gRNA core sequence followed by the BbsI cloning cassette and a rice tRNA^Gly^; and Cpf1gRNAPcr2fwd *ACAGACCCGGGTTCGATTCCCGGCTGGTGCATTTTTTGCCTACCTGGAGCCTGAGA* and Cpf1gRNAPcr2rev *TATACTGTTGCCGAGCACAATTGTCTAGAATGCATACGCATTAAGCGAAC* to generate a fragment encoding the genomic U6:3 terminator sequence directly downstream of rice tRNA^Gly^. *pCFD7* is available from Addgene (Plasmid 73916).

### Cas9 and Cpf1 expression plasmids

#### UAS-cas9.P2

We previously found that expressing high levels of Cas9 with the *Gal4/UAS* system resulted in toxicity in *Drosophila*, and that this was independent of endonuclease activity (Port et al., 2014). To attempt to reduce toxicity, we generated *UAS-cas9*.*P2*. This vector was designed to express lower levels of Cas9 compared to our previous *UAS-cas9*.*P1* plasmid (Port et al., 2014). To this end we used a *cas9* codon-optimised for expression in human cells (gift from George Church (Mali et al., 2013b), Addgene 41815) instead of the fly codon-optimised *cas9* present in *UAS-cas9*.*P1*. We also used the *pBIC-UASC* backbone (gift from Brian McCabe (Wang et al., 2012), Addgene 35200) instead of *pJFRC081* (gift from Gerald Rubin (Pfeiffer et al., 2012), Addgene 36432), as the latter plasmid is optimised for high protein expression levels. *pBIC-UASC* was digested with EcoRI and XbaI and gel purified. *hCas9* was amplified from plasmid *act-cas9* (Port et al., 2014) with primers UAScas9P2fwd *CAACTTTGAATCACAAGACGCATACCAAACGAATTCATGGACAAGAAGTACTCCATTG* and UAScas9P2rev *AGAAGTAAGGTTCCTTCACAAAGATCCTCTAGATCACACCTTCCTCTTCTTCTTGGG*. The backbone and insert were assembled by Gibson Assembly. Toxicity was markedly reduced with *UAS-cas9*.*P2*. Whereas expression of a single copy of *UAS-cas9*.*P1* with *act-Gal4* or *hh-Gal4* resulted in lethality, flies with *act-Gal4* or *hh-Gal4* and a single copy of *UAS-cas9*.*P2* flies were viable and fertile. Some loss of posterior wing margin tissue was observed when two copies of *UAS-cas9*.*P2* were expressed from a *hh-Gal4* driver at 25°C, suggestive of residual toxicity. Plasmid *UAS-cas9*.*P2* should become available from Addgene in April or May 2016.

#### act-hAsCpf1

The plasmid *act-cas9* (Port et al., 2014) was digested with EcoRI and XhoI and gel purified. AsCpf1 was amplified from pY010 (gift from Feng Zhang (Zetsche et al., 2015), Addgene 69982) using primers AsCpf1fwd *GCTTACAGGATCGATCCCCGGGAATTCACCATGACACAGTTCGAGGGCTTTACC* and AsCpf1rev *TCACAAAGATCCTCTAGAGGTACCCTCGAGTTAGGCATAGTCGGGGACATCAT*. The backbone and insert were assembled by Gibson Assembly. *act-hAsCpf1* is available from Addgene (Plasmid 73917).

### Fly transgenesis and culture

In all experiments, transgenes were present in one copy. Transgenic fly lines were generated by standard procedures (Port et al., 2014). All transgenes used in this study were integrated at the same genomic landing site (P{y[+t7.7]CaryP}attP40) on the second chromosome using the PhiC31/attP/attB system. The use of a single landing site allows direct comparison of the activity of different gRNAs. A second independent integration of *UAS-cas9*.*P2* was generated at (P{y[+t7.7]CaryP}attP2) on the third chromosome. All crosses were performed at 25°C with 50 ± 5% relative humidity and a 12-h light/12-h dark cycle. Virgin females transgenic for Cas9 or Cpf1 were crossed to males expressing transgenic gRNAs. *UAS-cas9*.*P2* flies are available from the Bloomington Stock Center (Stock numbers 58985 and 58986) and *act-cpf1* flies should become available there in April or May 2016. Additional fly strains used in this study are presented in Supplementary Table 2.

### Evaluating mutagenesis rates

For the data in Fig. 1b and 3b, germline transmission of non-functional CRISPR alleles was evaluated by crosses to *e* or *se* homozygous mutant adults (the *se* mutant allele (*se^Δ5^*) was generated in a previous series of CRISPR experiments (F. Port, unpublished)). Progeny with ebony or sepia pigmentation phenotypes indicated transmission of a non-functional CRISPR allele. In other cases, germline transmission of CRISPR-induced mutations was evaluated by extraction of genomic DNA from progeny of *act-cas9/gRNA* flies x OR-R wild-type flies, PCR of the target site and Sanger sequencing (Port et al., 2014). Heterozygosity for indels could be reliably determined from mixed sequencing traces near the target site (Port et al., 2014). Evaluation of somatic targeting of *se* in flies (Fig. 1c) was performed as described (Port et al., 2015).

### Immunohistochemistry

*Drosophila* wing imaginal discs were dissected from third instar larva in ice cold PBS and immediately fixed in 4% formaldehyde in PBT (PBS containing 0.3% Triton-X 100) for 25 min at room temperature. Discs were washed three times in PBT for 10 min each at room temperature and then incubated in primary antibody (rabbit anti-Wls 1:1000, (Port et al., 2008)) in PBT at 4°C overnight. Discs were then washed three times (20 min, room temperature) in PBT containing 1% heat-inactivated goat serum, followed by incubation with secondary antibody (Alexa Fluor 555 goat anti-rabbit (Invitogen), 1:500 in PBT) (2 h, room temperature). Samples were washed three times in PBT and once in PBS before mounting in Vectashield containing DAPI (Vector Laboratories). Double-sided tape was used as a spacer between the microscopy slide and the coverslip to avoid compression of the wing discs.

### Image Acquisition

Wing imaginal discs were imaged on a Zeiss LSM 780 laser-scanning confocal microscope using the sequential scanning mode and a 40x/1.3NA oil objective. Adult wings were mounted in 50% glycerol / 50% ethanol and imaged on a Nikon Eclipse TS100 microscope with a 5x objective and a Nikon Coolpix digital camera.

### Image Analysis

Image analysis was performed in Fiji (Schindelin et al., 2012). Brightness and contrast was adjusted to a comparable extent in all images in one series. The “measure’ tool was used to determine the areas of wing imaginal discs and adult wings that displayed the phenotypes documented in Fig. 2c, d and f.

## AUTHOR CONTRIBUTIONS

F.P. conceived the study, designed experiments, performed experiments, analysed data and wrote the manuscript. S.B. designed experiments, analysed data and wrote the manuscript.

## ACKNOWLEDGEMENTS

We would like to thank Nadine Muschalik for extensive input and discussions, as well as other members of the Bullock lab and users of www.crisprflydesign.org for feedback. This study was supported by a Marie-Curie IntraEuropean Fellowship (to F.P.) and a UK Medical Research Council Project U105178790 (to S.B.).

